# Functional divergence of TBP homologs through distinct DNA binding dynamics

**DOI:** 10.1101/2024.11.07.622486

**Authors:** Jieying H. Cui, James Z.J. Kwan, Armin Faghihi, Thomas F. Nguyen, Sheila S. Teves

## Abstract

The TATA box-binding protein (TBP) is an evolutionarily conserved basal transcription factor common in the pre-initiation complex of all three eukaryotic RNA polymerases (RNA Pols). Despite their high conservation, homologous TBPs exhibit species- and tissue-specific functions that may contribute to the increasingly complex gene expression regulation across evolutionary time. To determine the molecular mechanisms of species- and tissue-specificity for homologous TBPs, we examined the ability of yeast TBP and murine TBP paralogs to replace the endogenous TBP in mouse embryonic stem cells. We show that, despite the high conservation in the DNA binding domain among the homologs, they cannot fully rescue the lethality of TBP depletion in mESCs, largely due to their inability to support RNA Pol III transcription. Furthermore, we show that the homologs differentially support stress-induced transcription reprogramming, with the divergent N-terminal domain playing a role in modulating changes in transcriptional response. Lastly, we show that the homologs have vastly different DNA binding dynamics, suggesting a potential mechanism for the distinct functional behavior observed among the homologs. Taken together, these data show a remarkable balance between flexibility and essentiality for the different functions of homologous TBP in eukaryotic transcription.

## Introduction

Transcription initiation is the first step in gene expression, and requires the hierarchical assembly of the RNA Polymerase (RNA Pol) machinery at the promoters of genes (1, 2). Although the three main eukaryotic RNA Pols require general transcription factors that are specific to each system, the TATA-box binding protein (TBP) remains uniquely common to all three (3–7). TBP, as part of TFIID, is believed to be the first factor to bind onto the promoter region and trigger the hierarchical assembly of the other factors (8, 9). TBP consists of an unstructured N-terminal domain (NTD) and a conserved DNA binding core that recognizes the TATA-box DNA binding motif (10–12). This highly conserved transcription factor is present throughout archaea and eukarya (13), and has been shown to be essential for viability in all species.

Throughout evolution, gene duplication and horizontal gene transfer events have led to the emergence of TBP paralogs, enabling increasingly complex gene regulatory mechanisms (14, 15). The three main eukaryotic TBP paralogs are known as TBP-related factors 1-3 (TRF1, TRF2 and TRF3) (16–18). TRF1 is found primarily in insects (16), whereas metazoan-specific TRF2 and vertebrate-specific TRF3 are found broadly throughout multicellular organisms with tissue-specific expression patterns (17, 18). For example, in mice and humans, TBP and TRF2 are broadly expressed in many cell types, but TRF2 is highly expressed in spermatocytes and TRF3 expression is largely restricted to oocytes in mice (19–22). Furthermore, both paralogs play important roles in reproductive systems, with TRF2 and TRF3 knockout in mice leading to male and female sterility, respectively (23–25).

Despite the emergence of paralogs, TBP remains the predominant initiation factor. In vitro reconstitution of mammalian transcription by each of the three RNA Pols requires TBP for proper initiation (26, 27). However, recent studies in mouse and human cells suggest that in vivo transcription by RNA Pol I and II have relaxed their requirement for TBP (28–30). Specifically, these studies showed that rapid degradation of TBP in mouse embryonic stem cells (mESCs) and human HAP1 cells does not affect RNA Pol I and II activity but leads to massive downregulation of RNA Pol III transcription (28–30). In contrast, TBP remains essential for all transcription in yeast cells (4, 6). One potential mechanism to compensate for the species-specificity of TBP is the emergence of the TBP paralogs, which is absent in unicellular eukaryotes such as yeast. However, the tissue-specificity of the paralogs suggests that a functional divergence has also occurred between yeast and mammalian TBP.

In this study, we assessed how the conservation of TBP homologs leads to diverse species- and tissue-specific functions. Using an acute depletion system for TBP in mESCs, we examined if different TBP homologs, specifically yeast TBP and the mouse paralogs TRF2 and TRF3, can functionally replace the endogenous TBP protein. Despite the varying degrees of conservation in the DNA binding domain, TBP homologs cannot fully rescue mESCs viability upon TBP depletion. Each homolog also demonstrated a unique DNA binding profile, with TRF2 but not TRF3 partially rescuing RNA Pol III transcription upon depletion of TBP. Furthermore, we demonstrated that the divergent N-terminal domain of the homologs plays a specific role in fine-tuning changes in response to stress-induced transcriptional reprogramming. Lastly, we observed inherent differences in DNA binding dynamics for each TBP homolog, pointing to an evolved property that may enable species- and tissue-specific molecular behaviors, despite their high sequence conservation.

## Materials and Methods

### Cell lines

For all cell lines used in this study, the parental line is JM8.N4 mouse ESCs, purchased from KOMP repository, RRID: CVCL_J962. C64 and C94 is a CRISPR-Cas9 genetically modified JM8.N4 cell line containing mAID-TBP knock-in obtained as previously described (31). C41 is a CRISPR-Cas9 modified cell line with a Halo-mTBP knock-in obtained as previously described (31). B8 is a CRISPR-Cas9 genetically modified C64 cell line with a *Tbpl1* complete gene knock-out as previously described (29). F6 is a CRISPR-Cas9 genetically modified C94 cell line with a *Tbpl1* complete gene knock-out. Generation of the overexpressed TRF2-HA cell line was previously described (29). The cell lines have been authenticated by STR profiling, and tested negative for mycoplasma contamination.

### Generation of the overexpressed HA or Halo tagged constructs cell line

The mESCs were grown to 50% confluency and transfected using 1 μg of the construct (HA-mTBP, HA-yTBP, TRF3-HA, TRF3c-HA, HA-mTBPc, HA-yNTD-mTBPc, Halo-yTBP, Halo-TRF2, Halo-TRF3, Halo-TRF3c, Halo-mTBPc, Halo-yNTD-mTBPc) together with 1 μg of the Super Piggybac transposase plasmid using Lipofectamine 2000 (Invitrogen 11668-019) according to the manufacturer’s protocol. After 24 hr, 500 μg/mL of G418 (Fisher BP673-5) was added to select for successful random integration of the construct. Cell media containing antibiotics was refreshed daily until all negative control cells were dead.

### Cell culture

The mESCs were cultured on 0.1% gelatin-coated plates in ESC media KnockOut DMEM (Corning) with 15% FBS (HyClone), 0.1 mM MEM non-essential amino acids (Gibco), 2 mM GlutaMAX (Gibco), 0.1 mM 2-mercaptoethanol (Sigma-Aldrich) and 1000 units/ml of ESGRO (Chem-icon). ES cells were fed daily, cultured at 37°C in a 5% CO2 incubator, and passaged every 2 days by trypsinization. For TRF2-HA and TRF3-HA overexpressed cell lines, TBP degradation was performed by addition of IAA at 500 μM final concentration, or DMSO as carrier control, to a confluent plate of cells for 6 or 24 hours. Heat shock was performed at 42°C in a 5% CO2 incubator for 30 minutes. For HS and IAA treatment, cells were first treated with 24 hours of IAA followed by an additional 30 minutes before being collected. All cell lines with Halo-tagged constructs were treated with 6 hours of IAA before live-cell imaging.

### Cell counting Kit-8 (CCK-8) assay

Cells were seeded in a 96-well plate at a density of 4000 cells per well, with 100 μl of culture medium. Following incubation for 24 hours, either DMSO (vehicle control) or IAA were added to the culture. After 0, 24 and 48 hours of treatment, cells were washed with PBS and added 100 μl media with 10% of CCK-8 solution (Ab228554, Abcam). The plate was then incubated in a light-restricted setting at 37°C for 2 hours. The absorbance was measured at 450 nm by a SUNRISE plate reader.

### Antibodies for Western blot

Primary antibodies: α-HA 1:5000 (EpiCypher 13-2010), α-Tubulin 1:7000 (Abcam ab6046), α-TAF4 1:5000 (Santa Cruz sc-136093), α-BRF1 1:1000 (Santa Cruz sc-390821), α-H3.3 1:5000 (Abnova H3F3B M01), α-Halo 1:3000 (Promega G9211), α-Flag 1:5000 (Sigma M2) α-TBP 1:3000 (mAbcam51841). Secondary antibodies: IRDye 800CW Goat anti-mouse (Licor 926-32210), IRDye 800CW Goat anti-rabbit (Licor 925-32211), IRDye 680RD Goat anti-mouse (Licor 926-68070), or IRDye 680RD Goat anti-rabbit (Licor 926-68701).

### Protein Sequence Alignment Analysis

Protein sequence alignment was done using UniProt with the following Uniprot IDs: mTBP (P29037), TRF2 (P62340), TRF3 (Q6SJ95), yTBP (P13393). For the core regions of mTBP, TRF3 and yTBP, amino acid sequences starting with MSGIVP until the end (∼181aa) were used. Alignment percentage for both full-length and core protein sequences and phylogenetic tree modeling was performed using the UniProt multiprotein alignment suite.

### Protein Structure Conservation Analysis

Protein sequence conservation modeling was performed using iCn3D, a web-based 3D structure viewer that allows for overlaying protein structures (32). Using the “align protein complexes” function, Alphafold uniprot IDs were inputted, structures were overlaid and coloured according to conserved versus non-conserved amino acid residues and RMSD and TM-scores were calculated. Only the aligned region (core domain) is shown as the NTD is variable in sequence and length.

### Protein Structure - DNA Interaction Analysis

Protein structure DNA interaction analysis was performed using Alphafold 3 (33). Full-length and core-only protein sequences for the homologs were used as described above. Using previously described TBP CUT&Tag (29), TBP peak centers were defined for three RNA POL II gene promoters: *Gapdh*, with a full TATA-box motif, *Agbl5*, which contains only the 4 bp TATA sequence (TATA-like), and *UBB*, which contained no TATA motif. DNA sequences used were as follows; Gapdh-TATA (CCCCCCACCATCCGGGTTCC**TATAAAT**ACGGACTGCAGCCCTCCCTGGT), Agbl5-TATA-like (GGTAGAGCGGAGGACTGTAGTCAGTA**CAATATGG**TAATCCTTAGGTCGCTG), and UBB-TATA-less (CACCAATCAGCGTCGGCCTCAT**CTTTGACT**CCTCACCAATCAGCGCTGGCGCC). Bases bolded represent the approximate predicted binding site (DNA kink) visualized on ChimeraX based on Alphafold 3 prediction. The predicted template modeling (pTM) and the interface predicted template modeling (ipTM) scores were calculated from Alphafold 3, with default confidence cut-offs at 0.5 and 0.6, respectively. Alphafold 3 structures were then exported to ChimeraX for structural remodeling, alignment and coloring.

### CUT&Tag

CUT&Tag version 2 or version 4 was performed as described (Kaya-Okur et al., 2019), but with the following modifications. Cells were harvested at room temperature and 100,000 mESCs were used per sample. With the EpiCypher pAG-Tn5 (EpiCypher EP151117) or Ablab pA-Tn5, cryopreserved Drosophila S2 cells were spiked in at 10 or 20% to account for a spike-in control, during the cell pelleting step before washing. Antibodies used include RNA Pol II (Cell Signaling Technology D1G3K), RNA Pol III (Santa Cruz Biotechnology sc-21754), Rabbit anti-Mouse IgG (Abcam ab46540), α-H3K27me3 (Cell Signaling Technology C36B11), α-HA (EpiCypher 13-2010). Secondary antibodies used include Guinea Pig anti-Rabbit IgG (Antibodies-Online ABIN101961) and Rabbit anti-Mouse IgG (Abcam ab46540). Secondary antibody incubation times were 1 hour at room temperature. The commercial Epicypher pAG-Tn5 or Ablab pA-Tn5 was added at a final concentration of 1:20. Paired end sequencing was performed at the UBC Biomedical Research Centre using the NextSeq2000 50 cycles.

### CUT&Tag analysis

Reads were mapped on mm10 genome build using Bowtie2 with the following parameters: --no-unal --local --very-sensitive-local --no-discordant --no-mixed --contain --overlap –dovetail --phred33 –I 10 –X 9999. PCR duplicate reads were kept as these sites may represent real sites from adapter insertion from Tn5 as per recommendation from the Henikoff lab. A normalization factor was determined from *Drosophila* melanogaster (normalized to the control) alignment from Bowtie2 mapping and used to scale IAA-treated samples to control during generation of bigwig files. Downstream analyses, heatmaps, TSS plots, Gene plots, k-means clustering were performed using IGV, DeepTools, and BedTools suite (34, 35). ComputeMatrix from deeptools was done using binsize 10. Replicates were averaged using the wigtools and wigToBigWig, which averages the reads after normalizing treatment samples to the control samples of each run using Drosophila spike-in.

### DGE analysis using edgeR bioconductor

Scatter plots of differential gene expression (DGE) analysis were performed using the DGE tool from the edgeR bioconductor package. Starting subsampled bam files were used for all Pol II samples. Read counts were obtained using featureCounts command with paired end specificity for the entire gene and pseudogenes were removed (36). Reads were then imported into R Studio and analyzed using the DGE tool following the Bioconductor manual using default parameters (i.e. 5% FDR). First, DGEList was used to specify reads and gene names, followed by gene annotations (optional) then filtering and normalization by TMM (default option). Read counts below 20 were filtered using filterByExpr (min.count=20). Once reads were normalized, the design matrix was built and dispersion was estimated to determine biological variation. The command glmFit was used to determine DEG and the command topTags was used to show the top genes. Scatter plots were generated using the plotMD function and significant genes were obtained along with the raw values for downstream analyses such as heatmap and table generation.

### Scatter plot read counts analyses and filtering

Read counts for scatter plots were obtained from bam files using the bedtools multicov command and plotted to the promoters of genes (–250 bp to TSS or -250 to TES for HS) (34). Read counts were then normalized to the scaling factor, summed and plotted as is or log transformed before being plotted. Regression analysis was then performed to obtain the slope and r^2^ value.

### CoIP

Cells were grown to 90% confluency on a 15 cm gelatinized plate. After indicated treatments, cells were washed with 1× PBS, trypsinized and pelleted by centrifuging at 600 × g. Cell pellets were then lysed in 1 mL of Lysis Buffer (200 mM NaCl, 25 mM HEPES, 1 mM MgCl2, 0.2 mM EDTA, 0.5% NP-40, 1× Roche complete inhibitor, 0.2 mM PMSF, 1 mM benzamidine). Cell lysates were passed through a 25 G needle five times and incubated on an end-over-end rotator for 30 minutes at 4°C. Samples were spun at max speed at 4°C for 5 minutes. Supernatant was transferred to a new tube and the pellet was then digested using MNase (New England Biolabs M0297S) by nutating at 37°C for 30 minutes. Digested pellet was centrifuged at max speed and the new supernatant was added to the previous supernatant. Lysates were then pre-cleared using 20 µL/sample of Protein G Dynabeads and incubated on an end-over-end rotator for 2 hours at 4°C. Samples were then placed on a magnetic rack and aliquoted (10% vol/vol for input, 45% vol/vol for α-HA IP, 45% vol/vol for α-IgG IP). Pull-down was performed on the cell lysates by adding 1 µg of α-HA (Sigma Roche 12CA5) or mouse α-IgG (Santa Cruz Biotechnology sc-2025) incubating on an end-over-end rotator overnight at 4°C. The next day, Protein G Dynabeads were added to the mixture and incubated at 4 °C for 1 hour. Beads were washed six times with 600 µL Lysis Buffer. Proteins were eluted from the beads by addition of loading buffer and boiling at 100°C for 10 minutes.

### Single particle tracking: slow tracking

Cells were grown on gelatin-coated 35mm glass-bottomed dishes (Ibidi, μ-Dish 35 mm, high Glass Bottom 81158) at 37°C in a 5% CO2 incubator for 24 hours. Cells (HaloTagged TBPs) were labeled with 25pM JF549 (Lavis Lab) for 30 minutes at 37°C in a 5% CO2 incubator and washed 3 times with ESC media without phenol-red for 5 minutes each. Halo-TRF3 and Halo-TRF3c cell lines were dually labeled with 25nM JF646 after 25pM of JF549 labeling for epifluorescence imaging. Cells were imaged in ESC media without phenol-red. Imaging was conducted on a custom-built 3i (Intelligent Imaging Innovations) microscope equipped with an Alpha Plan-Apochromat 100×/1.46 NA oil-immersion TIRF M27 objective, EM-CCD camera (Andor iXon Ultra 897), a Zeiss Definite Focus 2 system and a motorized mirror to achieve HiLo-illumination. The customized laser launch includes 405 nm (350 mW), 488 nm (300 mW), 561 nm (1 W) and 640 nm (1 W) lasers. A multi-band dichroic (405 nm/488 nm/561 nm/633 nm quad-band bandpass filter) was used to reflect a 561 nm laser into the objective and emission light was filtered using a bandpass emission filter. The laser intensity was controlled using an acoustic-optic transmission filter (AOTF); a low constant laser intensity was used to minimize photobleaching. Images were collected at a frame rate of 2 Hz with 2% AOTF for a total of 1000 frames. Each Halo-tagged line was imaged in three biological replicates of 10–12 cells. Examples of raw SPT videos for H2B-Halo, Halo-mTBP, Halo-yTBP, Halo-TRF2 are shown in the Supplemental Data (Supplementary Videos 1–6). The Halo-TRF3-expressing mESCS were also imaged at a frame rate of 20 Hz and 30% AOTF for 400 frames to visualize more dynamic molecules and at 1% AOTF for epifluorescence imaging.

HaloTagged particles were identified using SLIMfast (37), a custom-written MATLAB implementation of the MTT algorithm (38), using the following algorithm settings: localization error: 10−6.25; exposure time: 200 ms; deflation loops: 3; number of gaps allowed: 1; maximum number of competitors: 5; maximal expected diffusion constant (μm^2^/s): 0.5. The residence times of HaloTagged molecules were determined using custom scripts as previously described (31, 39). Briefly, we quantified the dwell time of each HaloTagged molecule and generated a survival curve for bound HaloTagged TBPs as a function of time. A two-exponential function was fitted to the survival curve to determine apparent *k_off_*rates of HaloTagged TBPs in Fig. 6B-C. The photobleaching correction was performed by subtracting the apparent koff of H2B-Halo from the apparent *k_off_*of the HaloTagged TFs, and residence time was determined by taking the inverse of the photobleach-corrected koff for each HaloTagged TBPs.

### Single particle tracking: Fast tracking mode

Cells were grown and labeled as described in the previous section, and they were labeled with PA-JF549 dye to final 25 nM using the same condition as slow-tracking for each cell line. Imaging process for PA-JF549 dyes (40) performed by utilizing a multi-band dichroic filter (BrightLine quad-band bandpass filter, Semrock) to direct a 250 mW 549 nm laser (1 W, Coherent, Genesis) at 100% AOTF and 146 mW power of 405 nm laser (350 mW, Coherent, Obis) at 10% AOTF into the objective, while a bandpass emission filter (FF01 676/37 Semrock) was used to filter the emitted light. Fast-tracking imaging experiments involved capturing 10000 frames, with each experiment repeated three times for biological replicates and including 4 to 12 cells as technical replicates. At a frequency of 225 Hz, stroboscopic pulses of the 549 nm laser were applied for 1 ms per frame at maximum intensity to reduce motion blur. Single-molecule localization and tracking were conducted using SLIMfast, a MATLAB implementation of the MTT algorithm (38), with parameters and analysis methods applied as detailed previously (39), including settings for localization accuracy, competitor limits, and diffusion constant thresholds.

### Statistical analyses

Unless stated otherwise, P-values were calculated using a one-way ANOVA with a 95% confidence interval using GraphPad Prism suite.

### Immunofluorescence

Cells were grown on coverslips (Azn Scientific #ES0117650) that were pre-washed in 70% ethanol and coated with gelatin in tissue culture-treated six-well plates. After indicated treatments, cells were washed with 1 mL of PBS and fixed in 4% paraformaldehyde (UBC Chemical Stores #OR683105) for 15 min. After fixation, cells were washed and permeabilized with 1 mL 0.025% Triton-X for 5 minutes, followed by two 10-minute washes with PBS. Samples were blocked by nutating in PBS 5% BSA for 30 minutes. α-HA (Invitrogen SG77 715500) was diluted at 1:100 in PBS or α-Flag at 1:50 (Sigma M2), and added to cells in coverslips for 1 hour. Samples are washed twice with PBS (5 minutes for each wash). Coverslips were then incubated in Alexa Fluor 594 (A11005) 1:1000 for 1 hour, and washed with PBS twice (5 minutes for each wash). Samples are then incubated with DAPI (300 nM in PBS) for 5 minutes and washed twice with PBS (5 minutes for each wash). Coverslips were assembled using Vectashield mounting medium (BioLynx #VECTH1000). Fluorescent images were collected using the Leica DMI6000B inverted fluorescence microscope.

## Results

### Homologous TBPs do not fully rescue TBP lethality in mouse ESCs

Species- and tissue-specificities of the TBP homologs may arise from differences in non-conserved protein sequences. Multisequence alignment and phylogenetic analysis show that the TBP homologs (mouse and yeast TBP orthologs, and mouse paralogs TRF2 and TRF3) share 35%-65% similarity (Fig. S1A), but pairwise comparisons of the mTBP core (mTBPc) with TRF3 core (TRF3c), yeast TBP core (yTBPc), and TRF2, which lacks an NTD, shows 84%, 80%, and 41% similarity, respectively (Fig. 1A), and reflects the phylogenetic distance between homologs (Fig. 1B). Furthermore, pairwise comparisons of Alphafold2 predicted structures between the homologs and mTBP show root mean square deviation (RMSD) under 2Å and template-modeling score (TM score) over 0.9, suggesting highly similar structures among the homologs but with TRF2 as most different (Fig. 1C) (41). Using Alphafold3, we modeled the DNA binding capabilities of the homolog cores on promoter sequences containing the canonical TATA-box motif (TATAWAW), a TATA-like sequence (only TATA), or no motif (TATA-less), extracting the interface predicted template modeling (ipTM) score as a measure of model accuracy (33). All homologs, either full length or core only, show decreasing ipTM scores from TATA-box to TATAless promoters (Fig. 1D, Fig. S1B), but TRF2 shows the lowest ipTM scores for all promoter types, with some scores falling below the 0.6 cutoff for model confidence (Fig. 1D, Fig. S1B). These analyses suggest that mTBP, yTBP and TRF3 would function similarly in terms of binding DNA compared to TRF2.

**Figure 1.**
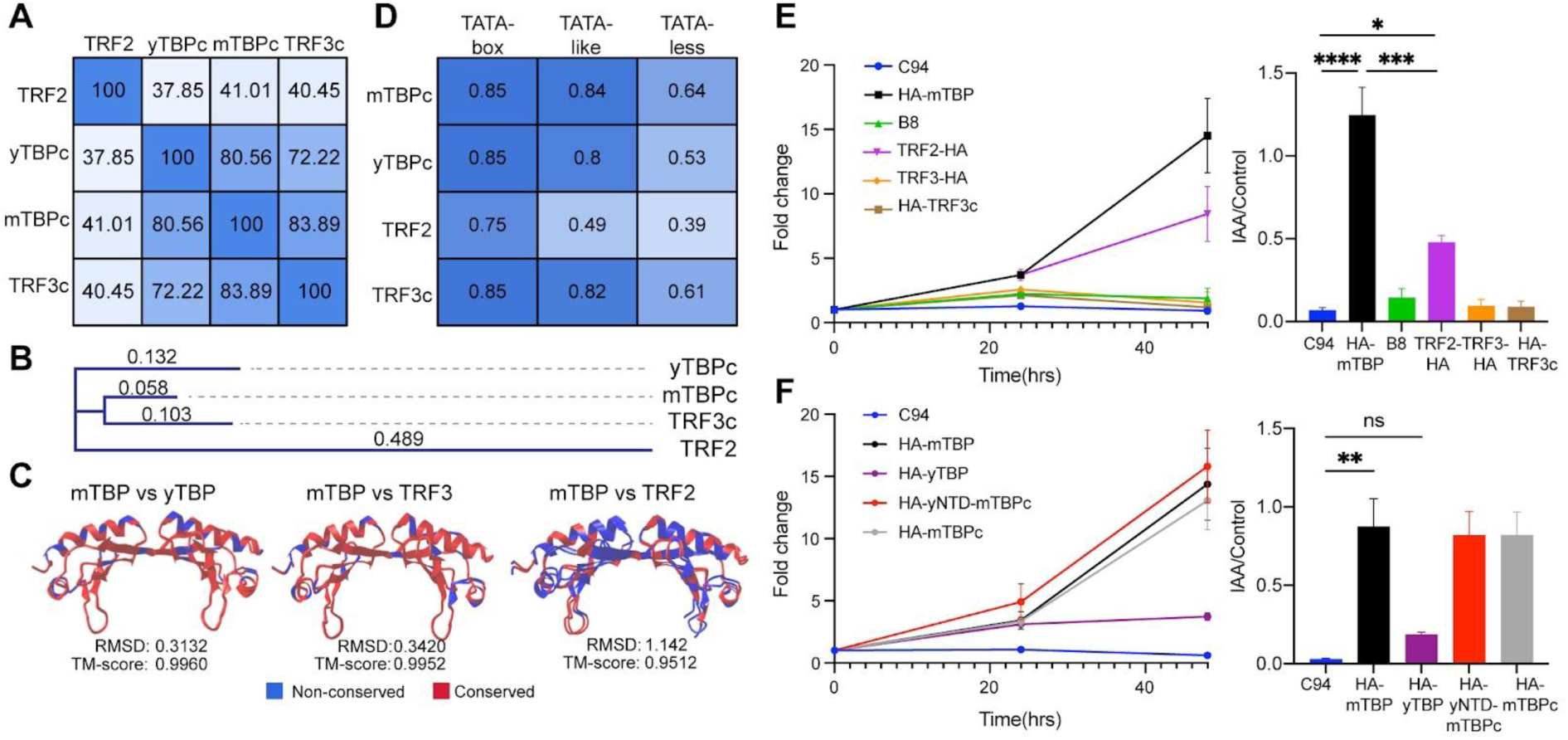
Conservation among the TATA box binding protein (TBP) homologs does not predict the ability to rescue lethality from TBP depletion. (A) Amino acid sequence percent similarity in the core domain of TBP homologs depicted as a heatmap matrix. (B) Evolutionary distance between TBP homologs based on conservation of the core domains. Branch lengths are indicated. (C) The structure of the homologs is modeled using iCn3D, utilizing Alphafold2 predicted protein structures and overlaying the protein structure conservation was analyzed using iCn3D as a pair-wise comparison against mTBP. RMSD, root mean square deviation; TM-score, template modeling score. (D) Alphafold3 was used to predict DNA binding of the core domains of the homologs to 3 types of promoters: TATA-box, TATA-like, and TATA-less. The ipTM scores, indicating the confidence in the predicted model, are represented as a matrix heatmap. (E) CCK8 growth curves (left) of parental mESCs (C94, B8) and mESCs expressing TRF2-HA, TRF3-HA and TRF3c-HA for 48 hours after IAA treatment to deplete endogenous TBP (right). The growth values at 48 hours for IAA treated mESCs are normalized to untreated control to calculate the percent rescue, and plotted as a bar graph with error bars representing standard error of the mean (SEM). (F) Same as E but for HA-mTBP, HA-yTBP, HA-yNTD-mTBP, and HA-mTBPc. Statistics calculated using one-way ANOVA: ns, non-significant; * p-value ≤ 0.05; ** p-value ≤ 0.01, **** p-value ≤ 0.0001.

To test this prediction, we designed a TBP replacement assay using a previously described homozygous knock-in of the minimal auxin inducible degron (mAID) to TBP (C94 mESCs), and subsequent knock-out of TRF2 (F6 mESCs). These mESCs do not express TRF3 (29). We stably integrated the HA-tagged wild-type mouse and yeast TBP (HA-mTBP, HA-yTBP) orthologs in C94 mESCs, and the HA-tagged TRF3 paralog in F6 mESCs. To test the role of the variable NTD, the HA-tagged truncated core of TRF3 (HA-TRF3c) was stably integrated in F6, and the HA-tagged mouse TBP core (HA-mTBPc) and a chimeric yeast NTD fused to mTBPc (HA-yNTD-mTBPc) were stably integrated in C94 individually (Fig. S1C). The TRF2-HA mESCs were generated and described previously (29). Expression of homologs were detected by Western blot, and nuclear localization was detected by immunofluorescence analysis, with TRF3 and TRF3c showing both nuclear and cytoplasmic signals (Fig. S1D-E). In each cell line, when the IAA drug is added, the endogenous mAID-TBP is rapidly degraded, enabling a functional test of cell viability for each homolog and fusion protein.

Using the CCK8 growth assay with and without IAA (Fig. 1E-F, Fig. S1F), we measured cell growth and viability of each homolog and calculated the ratio of growth at 48 hours for IAA over control to quantify percent rescue. As expected, IAA treatment of the parental cell line C94 resulted in ∼90% cell death by 48 hours (Fig. 1E-F). No homolog fully rescued cell viability upon TBP depletion, but TRF2-HA demonstrated a partial rescue at ∼50%, whereas HA-yTBP weakly supported ∼25% rescue, and TRF3-HA showed no rescue (Fig. 1E-F). Surprisingly, TRF3c-HA also failed to rescue the loss of endogenous TBP (Fig. 1E), despite being most similar to HA-mTBPc, which fully rescued cell viability along with the chimeric HA-yNTD-mTBPc (Fig. 1F). These results suggest that the core domain, not the variable NTD, dictates the ability to support cell viability, and that sequence and structural similarity between TBP homologs do not predict functional performance.

### Conservation in the core domain of TBP homologs does not predict DNA binding capability

To investigate the molecular mechanisms underlying the differences in cell viability, we measured the DNA binding capabilities of each homolog by performing CUT&Tag using the HA antibody targeting HA-mTBP, HA-yTBP and TRF3-HA in control and IAA-treated cells. We performed two independent replicates with spike-in normalization, and aligned the reads onto the mouse genome, with strong consistency between replicates (Fig. S2A-C). The TRF2-HA CUT&Tag was previously published (29) and the data was reanalyzed here. In both control and IAA-treated conditions, the wild-type HA-mTBP, TRF2-HA and HA-yTBP can recognize and bind onto the promoter of *Gapdh*, but TRF3-HA cannot (Fig. 2A). As previously described, TRF2-HA was bound to all transcribed RNA Pol II genes (29), but global binding somewhat decreased upon depletion of endogenous TBP (Fig. 2B), suggesting a potential synergy between these two paralogs. In contrast, TRF3-HA failed to bind onto RNA Pol II promoters (Fig. 2B). The ortholog HA-yTBP was bound to all expressed genes (Fig. 2B), but with about 44% times higher binding levels than HA-mTBP, as quantified by linear regression analysis (Fig. 2E, top).

**Figure 2.**
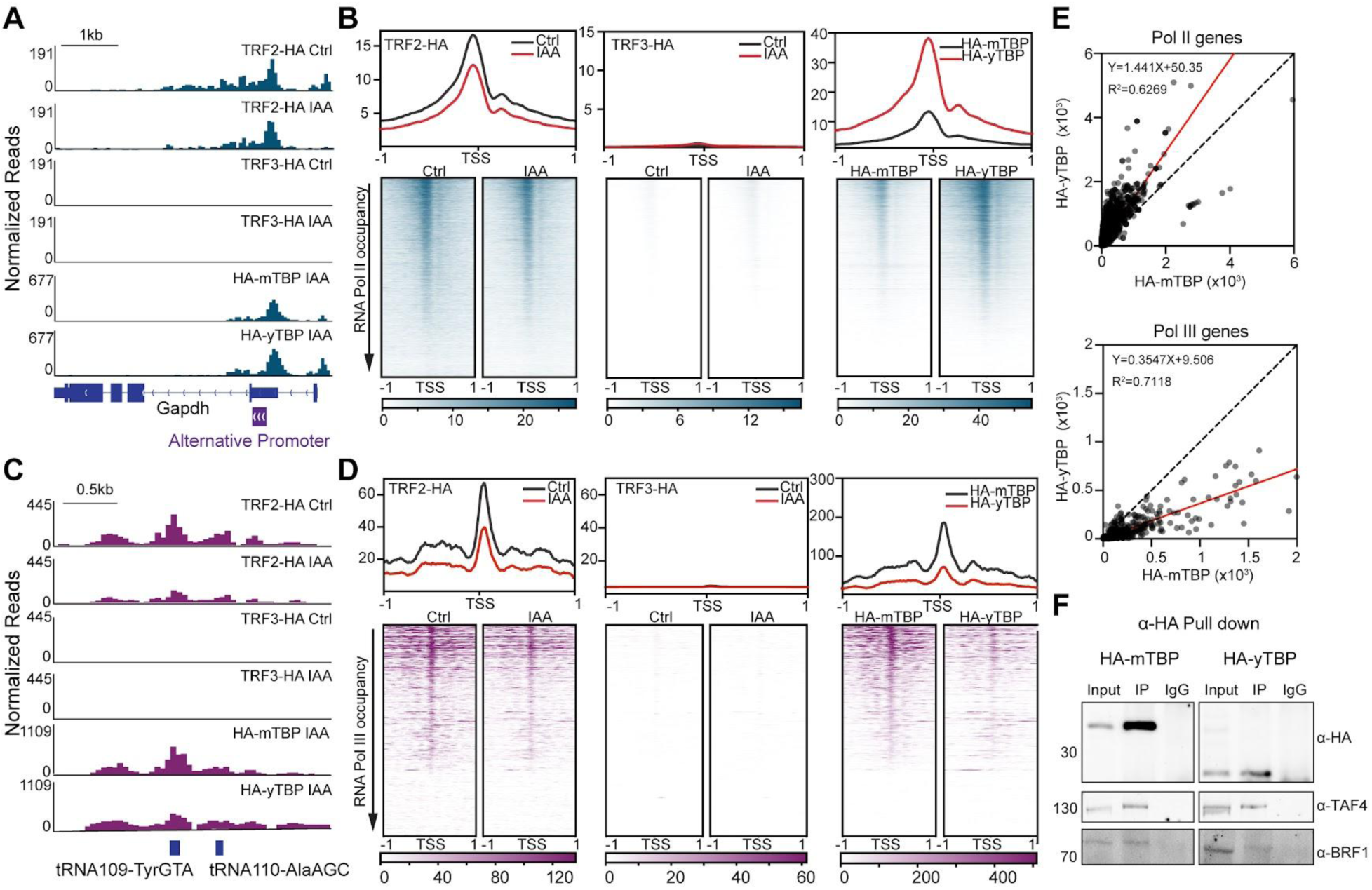
DNA binding of TBP homologs is independent of core domain conservation. (A) Gene browser tracks at *Gapdh* for ɑ-HA CUT&Tag of TRF2-HA, TRF3-HA, HA-mTBP and HA-yTBP in control (Ctrl, DMSO) or indole-3-acetic acid (500uM, IAA)-treated mESCs. Alternative promoters are indicated. (B) Genome-wide average plots (top) and heatmaps (bottom) arranged by decreasing Pol II occupancy of HA CUT&Tag for TRF2-HA (Ctrl vs IAA), TRF3-HA (Ctrl vs IAA), HA-mTBP (IAA) and HA-yTBP (IAA) in a 2 kb window surrounding the transcription start site (TSS) of all RNA Pol II genes. (C) Gene browser tracks at *tRNA109-TyrGTA* and *tRNA110-AlaAGC* for HA CUT&Tag of TRF2-HA, TRF3-HA, HA-mTBP and HA-yTBP in control or IAA-treated mESCs. (D) Genome-wide average plots (top) and heatmaps (bottom) arranged by decreasing Pol III occupancy of HA CUT&Tag for TRF2-HA (Ctrl vs IAA), TRF3-HA (Ctrl vs IAA), HA-mTBP (IAA) and HA-yTBP (IAA) in a 2 kb window surrounding the transcription start site (TSS) of all tRNA genes. (E) Normalized read counts of IAA-treated HA-yTBP CUT&Tag signal vs. IAA-treated HA-mTBP in the promoter (–250 to TSS) region of each Pol II (top) gene and in the gene body of each Pol III (bottom) gene are displayed as scatter plots. (F) Co-IP using ɑ-HA or IgG for cells expressing HA-mTBP (left) or HA-yTBP (right) and blotted with ɑ-HA, ɑ-TAF4, and ɑ-BRF1.

Unlike RNA Pol II, transcription by RNA Pol III in mESCs is critically dependent on TBP (29). At the *tRNA109-TryGTA* and *tRNA110-AlaAGC* loci, we observed that TRF2-HA binding is decreased when the endogenous TBP is depleted (Fig. 2C). The HA-yTBP exhibited a low level of binding whereas TRF3-HA showed no binding at these tRNA genes (Fig. 2C). Globally, TRF2-HA binding at tRNA genes decreased by over 50% when the endogenous TBP was depleted (Fig. 2D), consistent with the observed effect on RNA Pol II genes. TRF3-HA failed to bind to any tRNA genes (Fig. 2D). HA-yTBP showed some binding for all tRNA genes (Fig. 2D), albeit at 65% lower levels than the wild-type HA-mTBP as quantified by linear regression analysis (Fig. 2E, bottom), which is in stark contrast to the pattern at RNA Pol II promoters (Fig. 2B, E).

What determines the differences in DNA binding between the homologs, despite the strong similarity in protein sequence and structure? Our data show that HA-yTBP and TRF2-HA bound equally to gene promoters containing a defined TATA-box as to those without the conserved motif (Fig. S2D), consistent with previous studies showing sequence-independent binding of TBP in vivo (42, 43). For RNA Pol II initiation, structural studies have shown that mammalian TBP may be positioned by the TFIID complex (8), and in tRNA expression, the TFIIIC complex first binds the promoter and recruits the TBP-containing TFIIIB complex (44, 45). Using co-immunoprecipitation (co-IP) assays, we observed that HA-yTBP interacts with both mouse TAF4 and BRF1 to a similar degree as HA-mTBP (Fig. 2F), providing a potential mechanism for DNA binding for the ortholog. However, previous studies have shown that mouse TRF2 and TRF3 do not interact with TFIID and instead form their own complexes with TFIIA (46, 47), suggesting that the paralogs have evolved an alternative pathway to DNA binding than the TBP orthologs.

### Failure to fully rescue lethality is due to the inability to support Pol III transcription

Given the different DNA binding capabilities we observed, we questioned how the homologs affect the recruitment of the RNA Pols. We first assessed the chromatin occupancy of RNA Pol II through CUT&Tag with and without IAA in the presence of different homologs. Consistent with previous results, RNA Pol II occupancy was not affected by the degradation of TBP or the expression of different TBP homologs, with consistent biological replicates (Fig. S3A-B) (29).

Next, we examined potential changes to RNA Pol III recruitment. We performed RNA Pol III CUT&Tag with or without the endogenous TBP in two biological replicates and spike-in normalization, with good consistency between replicates (Fig. S3C). RNA Pol III recruitment to *tRNA109-TyrGTA* and *tRNA110-AlaAGC* genes (Fig. 3A) and to all tRNAs (Fig. 3B) was significantly impaired when the endogenous TBP was depleted, consistent with previous findings (29). However, when TRF2-HA and HA-yTBP were expressed in the absence of endogenous TBP, we observed a partial rescue of RNA Pol III occupancy at these loci (Fig. 3A), and at all tRNA genes (Fig. 3B) to about 34% and 32% relative to wild type mTBP (Fig. 3C, E), respectively, which correlates with the homologs’ ability to partially bind onto tRNA genes. TRF3-HA did not support RNA Pol III recruitment at these loci (Fig. 3A) and at all tRNA genes (Fig. 3B, D) upon TBP degradation. Notably, TRF2-HA and HA-yTBP supported different subsets of tRNAs, which is not correlated by the expression levels of the tRNAs (Fig. 3F). The binding ability of TBP homologs at tRNA genes correlates with the ability to support RNA Pol III binding, which reflects the varying ability to rescue lethality due to TBP depletion.

**Figure 3.**
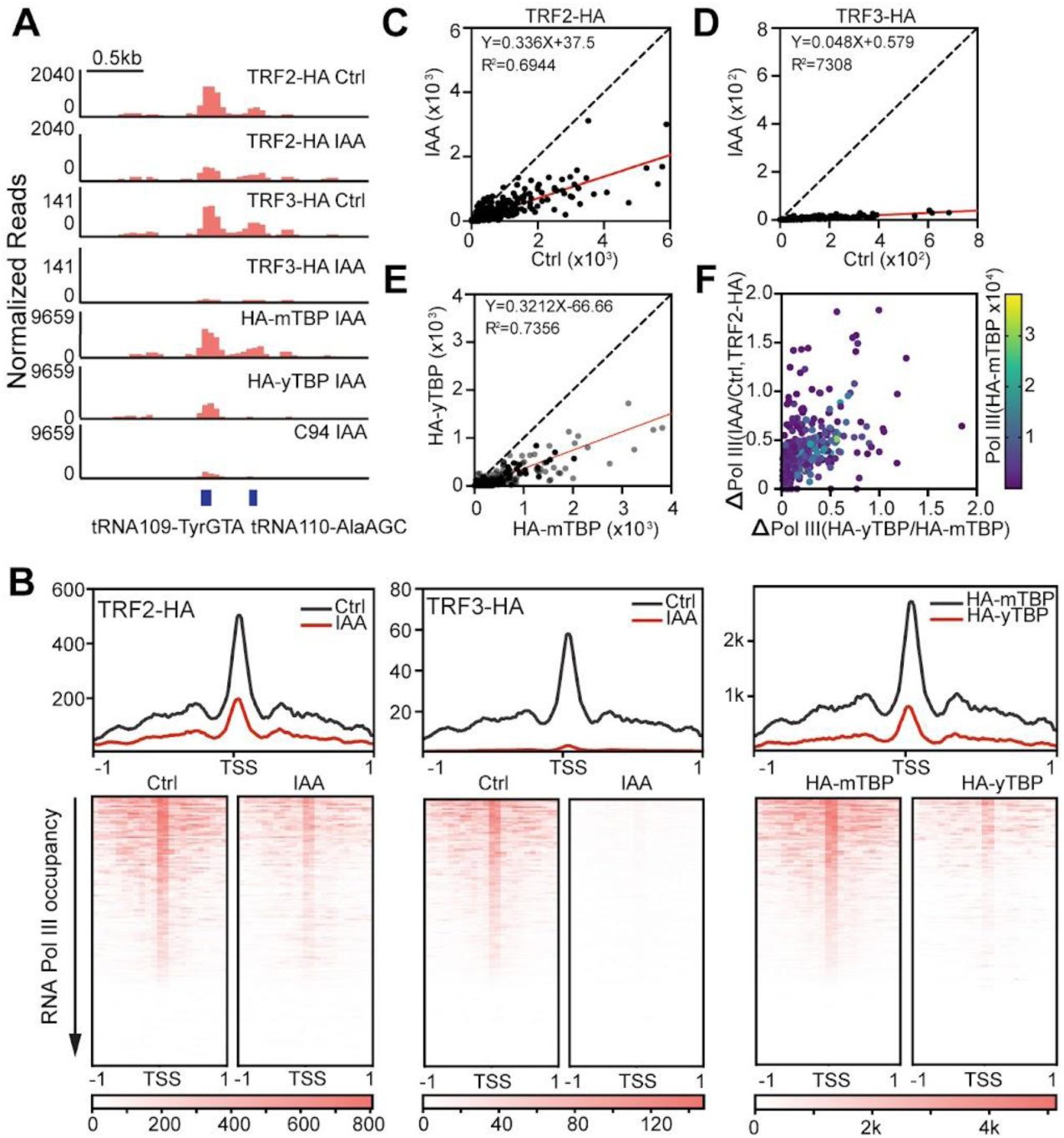
TBP homologs differentially affect RNA Pol III transcription of tRNA genes. (A) Gene browser tracks at *tRNA109-TyrGTA* and *tRNA110-AlaAGC* for Pol III CUT&Tag of mESCs expressing TRF2-HA, TRF3-HA, HA-mTBP and HA-yTBP in control or IAA-treated conditions. (B) Genome-wide average plots (top) and heatmaps (bottom) of Pol III CUT&Tag for mESCs expressing TRF2-HA (Ctl vs IAA), TRF3-HA (Ctl vs IAA), HA-mTBP (IAA) and HA-yTBP (IAA) in a 2 kb window surrounding the transcription start site (TSS) of all tRNA genes arranged by decreasing Pol III occupancy in control conditions. (C-E) Normalized read counts of Pol III CUT&Tag signal in TRF2-HA IAA vs Ctrl (C), TRF3-HA IAA vs Ctrl (D) and in IAA-treated HA-yTBP vs. IAA-treated HA-mTBP (E) in the gene body of each Pol III gene are displayed as scatter plots. (F) The change in the normalized read counts of Pol III CUT&Tag signal (IAA/Ctrl, TRF2-HA) vs the change in IAA treated Pol III CUT&Tag signal (HA-yTBP/HA-mTBP) are displayed as a scatter plot for each tRNA genes. The heatmap color represents the normalized read counts of Pol III CUT&Tag signal in IAA-treated HA-mTBP mESCs.

### The variable NTD does not affect DNA binding and function of TBP under homeostasis

The divergent NTD is variable both in sequence and length, with the NTDs for TRF3, mTBP, and yTBP at about 170, 130, and 60 amino acid residues, respectively (Fig. S1C). Although removal of the NTD in mTBP did not affect the growth phenotype of mESCs, we assessed its molecular role by performing CUT&Tag on the HA-mTBPc, TRF3c-HA, and the chimeric HA-yNTD-mTBPc, with and without IAA treatment. Biological replicates of CUT&Tag show strong consistency for each expressed homolog (Fig. S4B-C, E). At the *Gapdh* locus, the HA-mTBPc and HA-yNTD-mTBPc were bound to the promoter region at similar levels as the full length HA-mTBP, whereas TRF3c-HA failed to bind (Fig. 4A). This pattern is consistent for all RNA Pol II genes (Fig. S4A, D). Scatter plot analysis showed highly similar binding between the core and full length mTBP, whereas the HA-yNTD-mTBPc showed a 45% increase compared to HA-mTBP (Fig. 4B).

**Figure 4.**
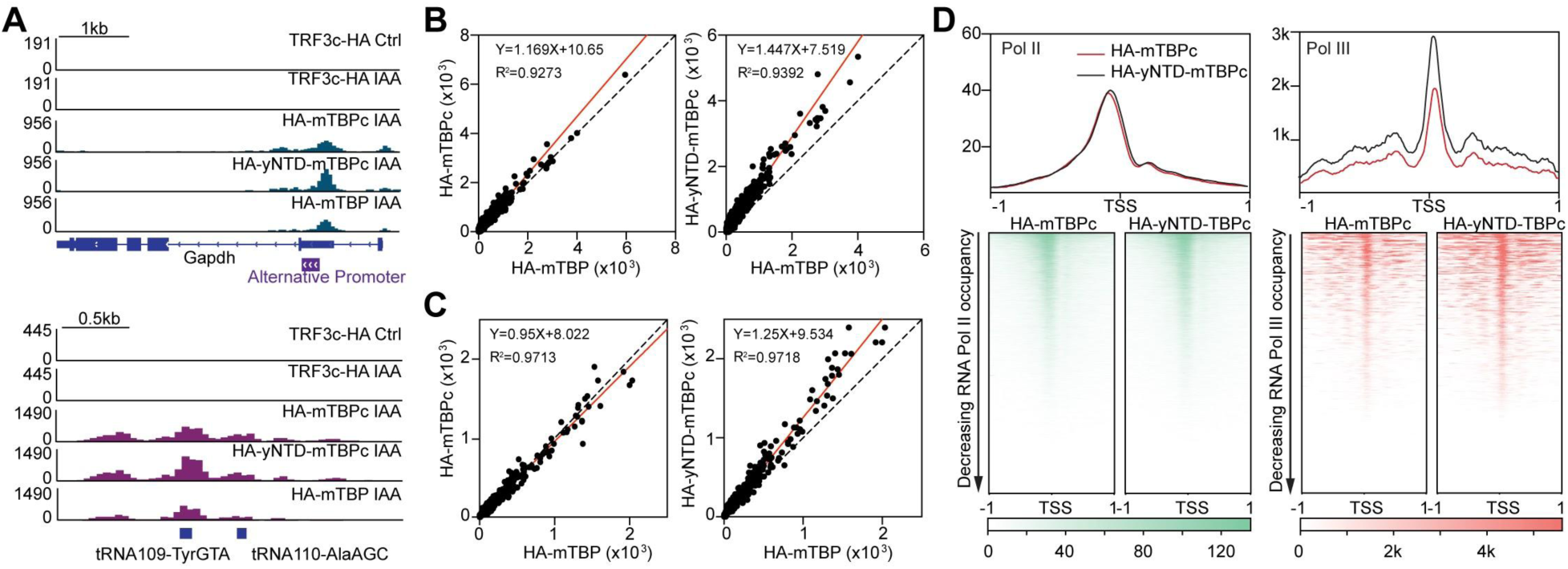
The variable NTD does not affect DNA binding and function of TBP under homeostasis. (A) Gene browser tracks at *Gapdh* for HA CUT&Tag of mESCs expressing TRF3c-HA, HA-mTBPc, HA-yNTD-mTBPc, HA-mTBP in control or IAA-treated conditions. Alternative promoters are indicated. (B) ɑ-HA CUT&Tag normalized read counts in the promoter region of each RNA Pol II gene for HA-mTBPc versus HA-mTBP (left) and HA-yNTD-mTBPc versus HA-mTBP (right) are displayed as scatter plots. (C) Gene browser tracks at *tRNA109-TyrGTA* and *tRNA110-AlaAGC* for HA CUT&Tag of mESCs expressing TRF3c-HA, HA-mTBPc, HA-yNTD-mTBPc, HA-mTBP in control or IAA-treated conditions. (D) tRNA gene body normalized read counts for HA CUT&TAg of HA-mTBPc versus HA-mTBP (left) and HA-yNTD-mTBPc versus HA-mTBP (right) are displayed as scatter plots. (E) Genome-wide average plots (top) and heatmaps (bottom) of CUT&TAg for RNA Pol II (left) and III (right) in mESCS expressing for HA-mTBPc, and HA-yNTD-mTBPc in a 2 kb window surrounding the transcription start site (TSS) of all RNA Pol II or tRNA genes.

We next examined the role of the NTD in homolog binding on tRNA genes. For mTBP, removal of the endogenous NTD or replacement with the yeast NTD showed no change in binding ability at the *tRNA109-TyrGTA* and *tRNA110-AlaAGC* genes (Fig. 4A, bottom). Heatmaps centered at tRNA TSSs (Fig. S4A, right) and scatter plot analysis (Fig. 4C) of HA-mTBPc showed highly similar binding levels at all tRNA genes compared to HA-mTBP, whereas HA-yNTD-mTBP showed 25% increase in binding compared to wild type. As with RNA Pol II genes, the removal of the NTD did not change the behavior of TRF3 binding on tRNA genes (Fig. 4A, S4D).

To measure effects on transcription, CUT&Tag of RNA Pol II and RNA Pol III was also performed for each cell line and plotted as heatmaps in a 2kb window around the TSS for all RNA Pol II genes or tRNA genes in two biological replicates (Fig. 4D, Fig. S4B-C). The RNA Pol II occupancy did not show significant changes in the truncated or chimeric products compared with either full-length protein, whereas the RNA Pol III had a minor increase in occupancy for the HA-yNTD-mTBPc, though there was no effect on cell viability (Fig. 4D, Fig. 1F). Taken together, we conclude that, although the highly divergent NTD may have minor effects on TBP binding at both RNA Pol II and III promoters, it has no major effects on RNA Pol II or III transcription at homeostasis.

### The variable NTD modulates transcriptional responses to stress

We next tested the role of the variable NTD under transcriptional reprogramming using the highly conserved heat shock (HS) response. Specifically, mESCs expressing HA-mTBP, HA-mTBPc, HA-yNTD-mTBPc, and HA-yTBP were exposed to 42°C for 30 minutes while the endogenous TBP was depleted, and CUT&Tag analysis for HA and RNA Pol II was performed in two biological replicates with spike-in normalization (Fig. S5D). Gene browser tracks for both HA and RNA Pol II CUT&Tag at the HS-induced gene *Hspa1a* showed massive increases in homolog and RNA Pol II binding upon HS compared to control samples at 37°C in all cell lines (Fig. 5A-B). DGE analysis of RNA Pol II CUT&Tag data in cells expressing HA-mTBP showed that 1679 and 1392 genes are significantly upregulated and downregulated, respectively, in HS samples compared to control (Fig. S5A).

**Figure 5.**
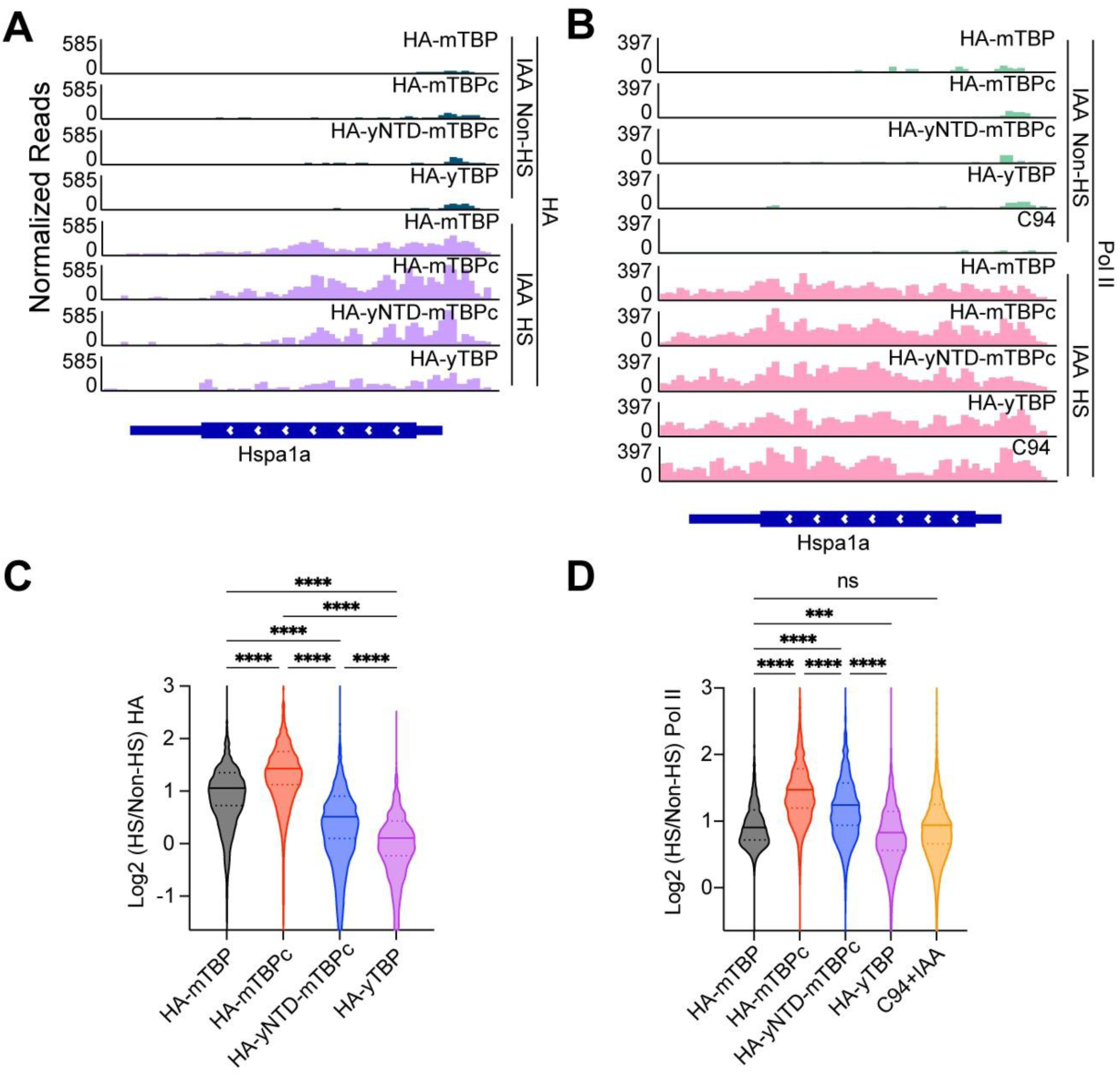
The variable NTD modulates transcriptional responses to stress. (A-B) Gene browser tracks at *Hspa1a* for HA (A) and Pol II (B) CUT&Tag of mESCs expressing HA-mTBP, HA-mTBPc, HA-yNTD-mTBPc, HA-yTBP in IAA-treated non-heat-shocked (Non-HS) or heat-shocked (HS) conditions. (C-D) The change in normalized read counts of HA (C) and Pol II (D) CUT&Tag signal in IAA-treated HA-mTBP, HA-mTBPc, HA-yNTD-mTBPc and HA-yTBP mESCs was quantified as Log 2 (HS/Non-HS) in the promoter and gene body of each Pol II gene, and visualized as a violin plot. Statistics calculated using one-way ANOVA: ns, non-significant; *** p-value ≤ 0.001; **** p-value ≤ 0.0001.

The change of induction was quantified by measuring the fold change in the promoter and gene body read counts of each upregulated gene at HS versus non-HS conditions. Although all HA-tagged homologs showed increased binding upon HS (Fig. S5B, D), the extent of this change varied among the expressed constructs. Notably, the truncated HA-mTBPc exhibited a greater increase in occupancy on HS-induced genes compared to the full-length HA-mTBP during HS (Fig. 5C). Additionally, incorporating the yeast NTD into mTBPc affected the chimeric TBP binding to HS-induced genes, resulting in a smaller increase in binding relative to HA-mTBPc and HA-mTBP (Fig. 5C). We also observed that HA-yTBP was less efficient in being recruited to HS-related genes compared with HA-mTBP in mESCs (Fig. 5C).

As previously observed, RNA Pol II in TBP-depleted C94 cells was still recruited to HS-induced genes (Fig. 5B), and is unaffected by the presence of exogenous HA-mTBP (Fig. 5B, D, S5C). Although gene induction does not require TBP, mESCs expressing the HA-mTBPc exhibited a greater change in RNA Pol II induction compared to cells expressing the full-length HA-mTBP (Fig. 5D), suggesting an exaggerated RNA Pol II response. This effect is somewhat modulated in the HA-yNTD-mTBPc cells, though the change in induction remains higher than the HA-mTBP (Fig. 5D). The HA-yTBP mESCs exhibited similar levels of RNA Pol II induction as the HA-mTBP and C94 samples, suggesting no modulating effects by the full-length ortholog (Fig. 5D). Taken together, these analyses show that, although the NTD of TBP plays a minor role during homeostasis, it may function as a modulator of RNA Pol II induction under transcriptional reprogramming during stress.

### TBP homologs display distinct DNA binding dynamics

We next examined the biophysical properties of each homolog by performing live-cell single molecule imaging and single particle tracking (SPT). First, we stably integrated Halo-tagged TBP versions in either the C94 (Halo-yTBP, Halo-mTBPc, Halo-yNTD-mTBPc), or F6 (Halo-TRF2, Halo TRF3, Halo-TRF3c) cell lines and measured expression by Western blot (Fig. S6A). For the Halo-mTBP, we used the previously described endogenous knock-in of HaloTag in the TBP locus (31).

To determine the residence times on DNA for each homolog, Halo-tagged expressing mESCs were labeled with Halo-specific JF549 and imaged in slow-tracking mode at 2 Hz (48), such that diffusing molecules ‘blur’ out while DNA-interacting molecules appear as diffraction-limited spots (Fig. 6A). Consistent with the CUT&Tag data, we observed diffraction-limited spots for Halo-mTBP, Halo-yTBP and Halo-TRF2, but not Halo-TRF3 (Fig. 6A), suggesting that Halo-TRF3 does not form stable interactions with DNA. Indeed, using a ten-fold faster frame rate (20 Hz), we were able to capture diffraction-limited spots for Halo-TRF3 (Fig. 6B), confirming much more dynamic DNA interactions compared to the other homologs. Using the SLIMfast algorithm (37), single molecules were localized and tracked, and the dwell times were plotted as semi-log histograms averaged across three biological replicates (Fig. 6C-D), with high consistency between replicates (Fig. S6B). A two-component exponential decay model was fitted to the dwell time curves, and the apparent residence times for each homolog was extracted and corrected for photobleaching as previously described (31, 39) (Fig. 6C-D). Consistent with previous studies, the average residence time of Halo-mTBP is about 80 seconds (Fig. 6C-D). Halo-yTBP and Halo-TRF2 showed decreased residence times compared to Halo-mTBP, at about 50s and 30s, respectively (Fig. 6C). Although we were able to detect diffraction-limited spots at faster frame rates, Halo-TRF3 was too dynamic to perform a similar analysis.

**Figure 6.**
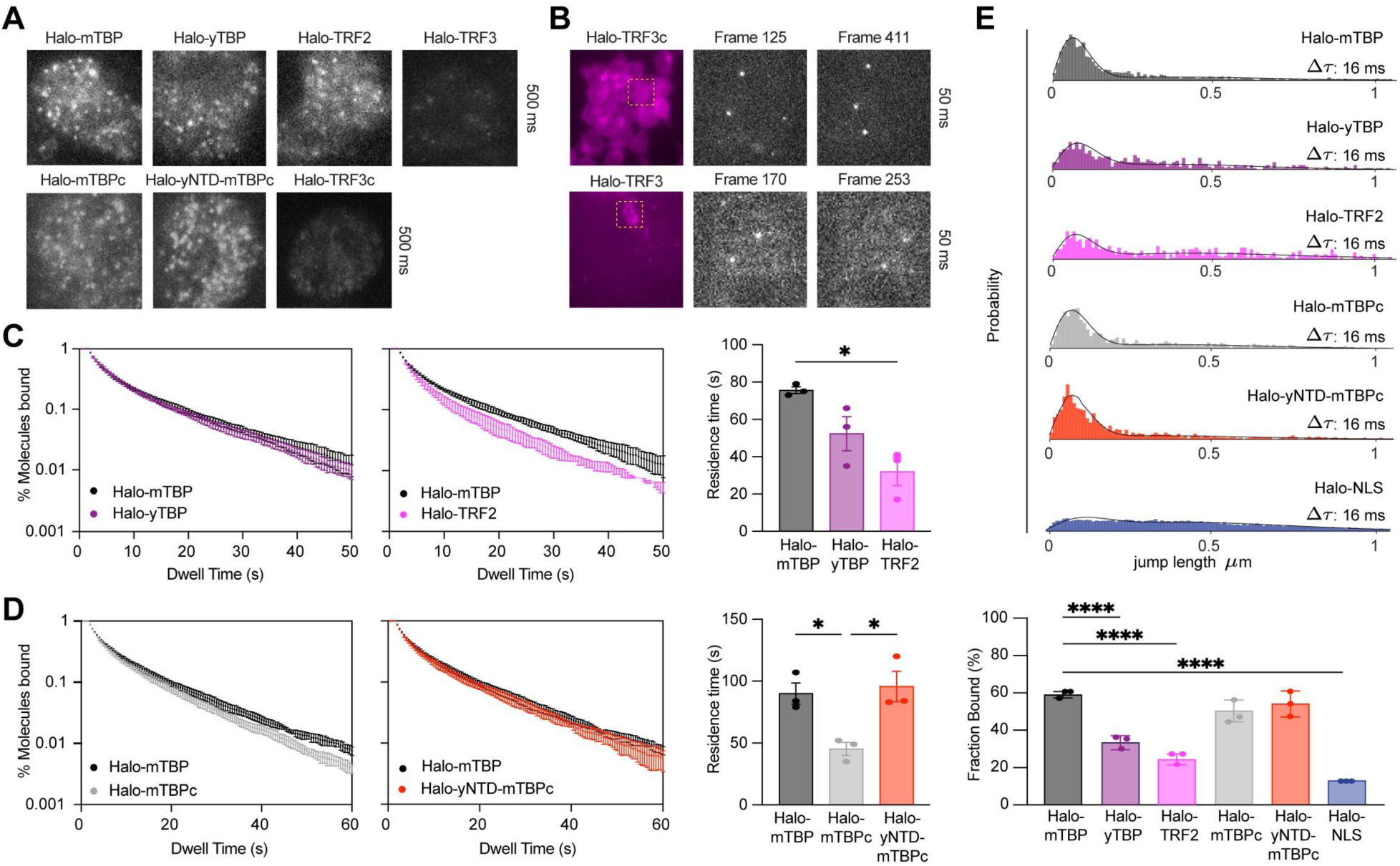
TBP homologs display distinct DNA binding dynamics. (A) Still images from SPT imaging of Halo-mTBP, Halo-yTBP, Halo-TRF2, Halo-TRF3, Halo-mTBPc, Halo-yNTD-mTBPc and Halo-TRF3c in IAA-treated mESCs indicating DNA-bound diffraction limited spots. (B) Epifluorescence and still images from SPT imaging of Halo-TRF3 and Halo-TRF3c using 20 Hz. (C-D) Dwell time histogram (left two panels) of the fraction of Halo-mTBP (C-D, black), Halo-yTBP (C, purple), Halo-TRF2 (C, pink), Halo-mTBPc (D, grey), Halo-yNTD-mTBPc (D, red) molecules remaining bound for IAA-treated mESCs. Quantification of the residence time (right) of each cell line. n = 3 biological replicates with 8-10 cells per replicate. Error bar represents SEM. (E) Jump length histogram measured in displacements (μm) after four consecutive frames (Δτ = 16 ms) during spaSPT for mESCs expressing Halotagged homologs or Halo-NLS as control (left). Fraction of bound molecules for each protein extracted from fitting the jump length histogram with a 2-state kinetic model. Error bar represents SEM. Statistics calculated using one-way ANOVA: ns, non-significant; * p-value ≤ 0.05; ** p-value ≤ 0.01, **** p-value ≤ 0.0001.

We assessed the contribution of NTD in DNA-binding residence times by performing slow tracking SPT for the truncation and chimeric proteins (Halo-TRF3c, Halo-mTBPc, Halo-yNTD-mTBPc) (Fig. S6B). Slow-tracking analysis shows that Halo-mTBPc is more dynamic than Halo-mTBP, with a residence time for the core at about half of the full-length TBP (Fig. 6D). Addition of the yNTD resulted in increased residence time for the chimeric protein to a level similar to the Halo-mTBP (Fig. 6D), suggesting that the NTD of both yeast and mouse TBP helps stabilize the mTBP core on chromatin (Fig. 6D). As with the full length Halo-TRF3, the DNA binding dynamics by the core-only Halo-TRF3c was detectable at 20 Hz but not at 2 Hz (Fig. 6A-B). These results suggest that DNA binding residence times are largely determined by the core domain, with some modulation by the NTD.

Next, we examined the dynamic behavior of the homologs. The mESCs expressing full length Halo-tagged TBP homologs were labeled with a photoactivatable dye (pa-JF549) and imaged at high illumination intensity using stroboscopic photoactivatable single particle tracking (spaSPT). Individual molecules were localized and their displacement between consecutive frames was measured as previously described (39, 49) and plotted as a histogram, with the HaloTag fused to a nuclear localization signal (Halo-NLS) as a non-DNA binding control (Fig. 6E, Fig. S6C). The higher proportion of short displacements (jump length) for Halo-mTBP suggests more DNA bound molecules compared to freely diffusing ones, as characterized by long displacements. In contrast, Halo-yTBP and Halo-TRF2 had smaller proportions of short displacements compared to Halo-mTBP, suggesting a higher population of freely diffusing molecules. Using Spot-On (49), we fitted a 2-state kinetic model on the displacement histograms to represent ‘bound’ and ‘free’ populations and the characteristic diffusion coefficients for each population. From these models, Halo-mTBP shows the highest proportion of bound molecules at 59%, whereas Halo-TRF2 and Halo-yTBP are at 24% and 33%, respectively (Fig. 6E). We also examined the diffusion coefficients of the freely diffusion molecules (Fig. S6D) but there are no statistically significant differences in diffusive behavior among the homologs. Since Halo-TRF3 did not exhibit measurable binding by SPT, we excluded this homolog from the analysis.

To test the role of NTD on TBP dynamics, we performed a similar analysis on the truncation (Halo-mTBPc) and chimeric (Halo-yNTD-mTBPc) constructs. Both versions exhibited binding percentages comparable to mouse TBP, with 50% for Halo-mTBPc (mouse TBP core) and 54% for Halo-yNTD-mTBPc (Fig. 6E). As with the homologs, these truncation and chimeric proteins display similar diffusive behaviors (Fig. S6D). Consistent with the slow-tracking results, these data suggest that the core region of TBP is the primary factor for determining the fraction of bound molecules and the dynamic behavior in the nucleus, whereas the NTD plays minor roles that may be important for fine-tuning changes in transcriptional responses.

## Discussion

In this study, we investigated the conserved and divergent properties of TBP homologs in mouse embryonic stem cells (mESCs). Despite high amino acid sequence conservation in the core domains, TBP homologs exhibited various capabilities in supporting cell viability, DNA binding, and transcription. Although mouse TRF3 shares the highest sequence similarity with mouse TBP in the core domain, it displayed the least conserved behavior at the functional and molecular level. In contrast, the more divergent yeast TBP and mouse TRF2 can partially support cell growth, DNA binding, and RNA Pol III transcription. Additionally, the highly variable N-terminal domain did not significantly impact DNA binding or ability to support RNA Pol II or III transcription in normal conditions, but influenced RNA Pol II transcriptional reprogramming during heat shock. Lastly, each homolog displayed distinct DNA binding dynamics that correlated with the functional and molecular phenotypes, suggesting that dynamics may be an evolved property of the core domain that can be partially modulated by the NTD.

It has been widely accepted that amino acid sequences determine protein structure, which in turn dictates biochemical function. As has been established in several studies (50, 51), protein pairs with sequence identity higher than 35–40% are very likely to be structurally similar. Here we provide an example where highly conserved protein sequences, with similar predicted structures, do not correlate well with their observed molecular and functional behaviors. How do the different TBP homologs exhibit such diverse molecular phenotypes despite high conservation in protein sequence and predicted structure? We propose that the distinct DNA-binding dynamics observed among the homologs provide an underlying mechanism for such functional diversity. In this model, subtle variations in DNA interaction and binding kinetics enable homologous TBPs to perform context-dependent functions, potentially affecting promoter recognition and recruitment of other general transcription factors. Our slow SPT data suggests that the majority of the differences in DNA-binding dynamics is largely due to key residues within the DNA-binding domain, with the NTD acting as a secondary modulating factor. As such, these dynamics could provide a level of regulatory specificity that underpins the distinct biological roles of TBP homologs in different species and tissues.

A secondary role for the NTD is somewhat surprising given that this sequence represents the largest divergence in TBP throughout evolution. Although deletion of the NTD in mice results in post-embryonic developmental lethality, yeast cells without the NTD are phenotypically indistinguishable from wild-type cells (52, 53). Yet at the molecular level, such species-specific differences in the NTD are minor and primarily manifest in altered transcriptional stress response (Fig. 4-5). It is possible that the effect of the NTD in DNA-binding dynamics may also be influenced by species- or tissue-specific cellular contexts such as differences in nuclear phase separation, which has been recognized as a mechanism for organizing nuclear components and modulating gene expression. Phase separation involves the formation of membrane-less condensates, where the transcriptional machinery, including TBP, may dynamically concentrate to regulate transcriptional processes. Evidence suggests that the human TBP may participate in phase separation, particularly in the context of disease where a poly-glutamine (poly-Q) stretch in the NTD is expanded (54). Intriguingly, this poly-Q stretch is conserved in the mouse TBP but absent in yeast NTD and the paralogs (15). In this context, the observed variations in DNA-binding kinetics attributed to the NTD (Fig. 6C) could influence their propensity to participate in or drive phase-separated condensates. For instance, TBPs with higher DNA-binding affinity and stability might be more likely to remain bound at promoter regions, potentially stabilizing transcriptional condensates. Conversely, truncated TBP with more transient DNA interactions might facilitate a more dynamic, fluid exchange of transcription factors within these condensates, thereby allowing for rapid transcriptional reprogramming under stress conditions. In this way, subtle changes in the conserved core can result in major changes in function, while large changes in NTD can have minor modulating effects, enabling a balance between flexibility and essentiality for TBP in eukaryotic transcription.

## Data, code, and materials availability

All sequencing data have been deposited in Gene Expression Omnibus (Accession number). All other data are available in the manuscript or in the supplementary materials.

## Supplementary Data

Supplementary videos are available separately.

## Funding

This work was supported by: The Canadian Institutes for Health Research Project Grant award to S.S.T. (PJT-162289); The National Sciences and Engineering Research Council Discovery Grant award to S.S.T. (RGPIN-2020-06106); The Stem Cell Network Early Career Researcher Jump Start Awards Program to S.S.T. (AWD-021244).

## Competing interests

Authors declare that they have no competing interests.

## Acknowledgments

We thank Dr. Eric Jan for providing S2 cells for spike-in normalization. We thank T. Stach (BRC-seq, UBC) for Illumina sequencing. This work was supported by Life Sciences Institute Cores (LSI Imaging, ubcFLOW, and QPCR Core), and by the UBC GREx Biological Resilience Initiative. For insightful comments on the manuscript, we thank Dr. Leann Howe. J.C. is supported by UBC 4-Year Fellowship, S.S.T is a Canada Research Chair and Michael Smith Foundation for Health Research Scholar.

## Author contribution

Conceptualization: JHC, JZJK, SST

Methodology: JHC, JZJK, AF, TFN

Investigation: JHC, JZJK, AF, TFN

Visualization: JHC, JZJK, AF, TFN, SST

Funding acquisition: JHC, SST

Supervision: SST

Writing – original draft: JHC, JZJK, AF, SST

Writing – review and editing: JHC, JZJK, TFN, SST

## Notes

### Competing Interest Statement

The authors have declared no competing interest.

