## Supplemental Figures for "Functional divergence of TBP homologs through distinct DNA binding dynamics"

### Supplementary Figure Legends

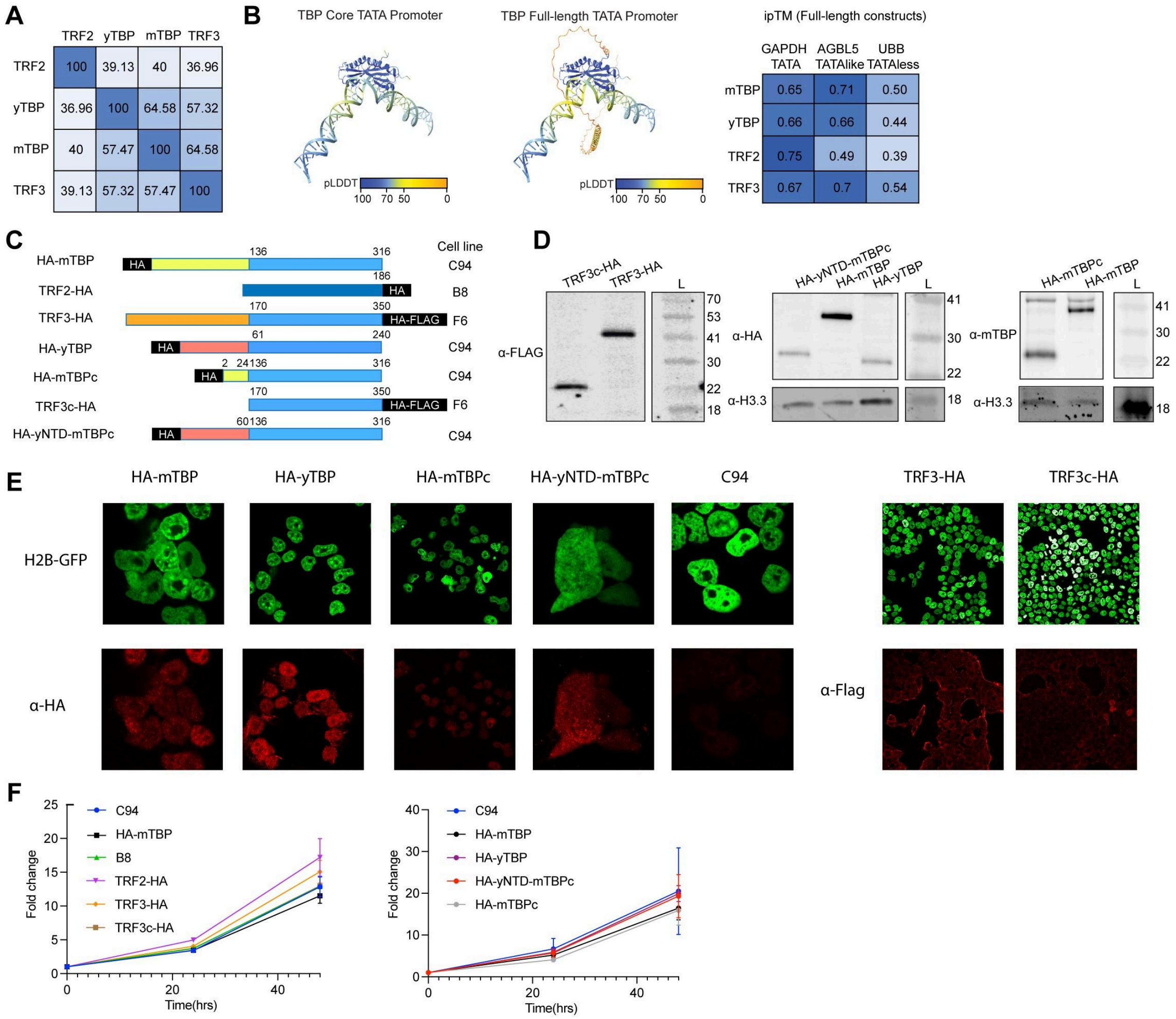


**Figure S1. Comparisons of TBP homologs and expression in mESCS.**

(A) Protein similarity heatmap matrix for full-length mTBP, yTBP, TRF2, and TRF3 as quantified by Uniprot multisequence alignment. (B) Predicted models for mTBPc (left) and TRF2 (middle) binding to the TATAbox-containing DNA sequence. pLDDT is the per-residue measure of local confidence ranging from 0 to 100 with higher scores indicating higher confidence and more accurate prediction. The ipTM scores for full length proteins and TATA-box, TATA-like, and TATA-less sequences are depicted as a heatmap matrix (right). (C) Depiction of each HA-tagged construct aligned at the core domain are shown along with the parental cell line that were used to overexpress the constructs. (D) Western blot analyses of each

HA-tagged homologs are shown. TRF3-HA and TRF3c-HA were detected using α-Flag; the HA-yNTD-mTBPc, HA-mTBP, and HA-yTBP were detected using α-HA, and the HA-mTBPc and HA-mTBP were confirmed using α-mTBP. Loading controls are depicted using α-H3.3. (E) Immunofluorescence analyses of each HA-tagged protein are shown. H2B-GFP was used to visualize the nucleus. HA-mTBP, HA-yTBP, HA-mTBPc, HA-yNTD-mTBPc and C94 were detected with α-HA and TRF3-HA and TRF3c-HA were detected using α-Flag. (F) CCK8 growth curves for each of the HA-tagged proteins are shown under DMSO-treated (no mAID-TBP depletion) conditions. Error bar represents SEM.

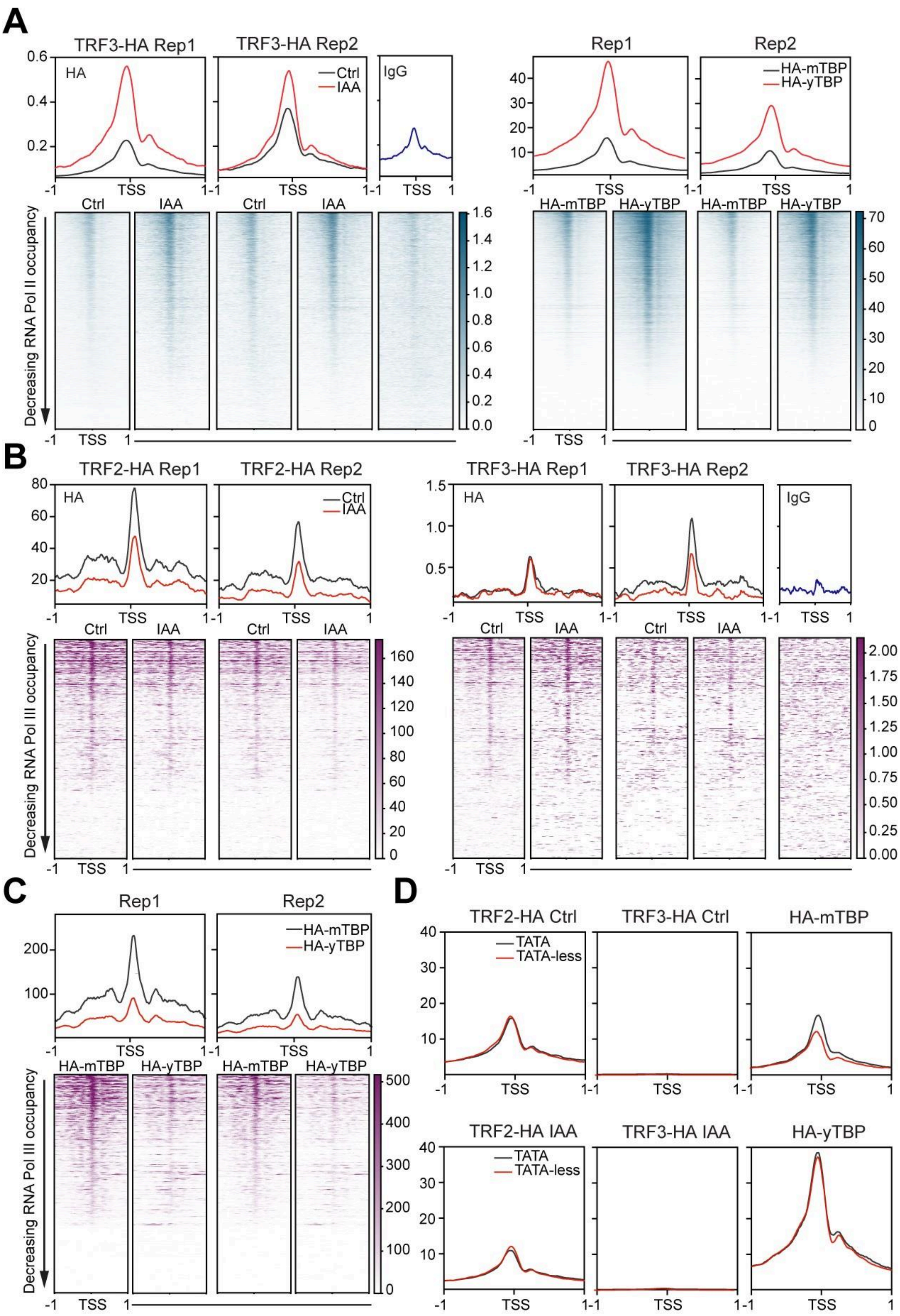

A
TRF3-HA Rep1
TRF3-HA Rep2
HA
Ctrl
IAA
IgG
TSS
Rep1
Rep2
HA-mTBP
HA-yTBP
Decreasing RNA Pol II occupancy
B
TRF2-HA Rep1
TRF2-HA Rep2
TRF3-HA Rep1
TRF3-HA Rep2
Decreasing RNA Pol III occupancy
C
Rep1
Rep2
HA-mTBP
HA-yTBP
Decreasing RNA Pol III occupancy
D
TRF2-HA Ctrl
TRF3-HA Ctrl
HA-mTBP
TRF2-HA IAA
TRF3-HA IAA
HA-yTBP
TATA
TATA-less

**Figure S2. Replicate analyses of TBP homologs' DNA binding at RNA Pol II and tRNA genes.**

(A) Biological replicates of HA CUT&Tag analyses displayed as average plots (top) and heatmaps (bottom) in a 2-kb window centered at the TSS of all RNA Pol II genes. Samples shown are TRF3-HA in DMSO- (control) and IAA-treated conditions, IgG (negative control for CUT&Tag, HA-mTBP and HA-yTBP. (B-C) Biological replicates of HA CUT&Tag analyses displayed as average plots (top) and heatmaps (bottom) in a 2-kb window centered at the TSS of all tRNA genes. Samples shown include TRF2-HA, TRF3-HA in DMSO- (control) and IAA-treated conditions, and IgG as negative control for CUT&Tag signal (B), and HA-mTBP and HA-yTBP (C). (D) Average plots of HA CUT&Tag signal for TATA-box containing (TATA) and TATA-less genes in a 2-kb window centered at the TSS. Samples analyzed include TRF2-HA and TRF3-HA (Control versus IAA-treated), HA-mTBP, and HA-yTBP.

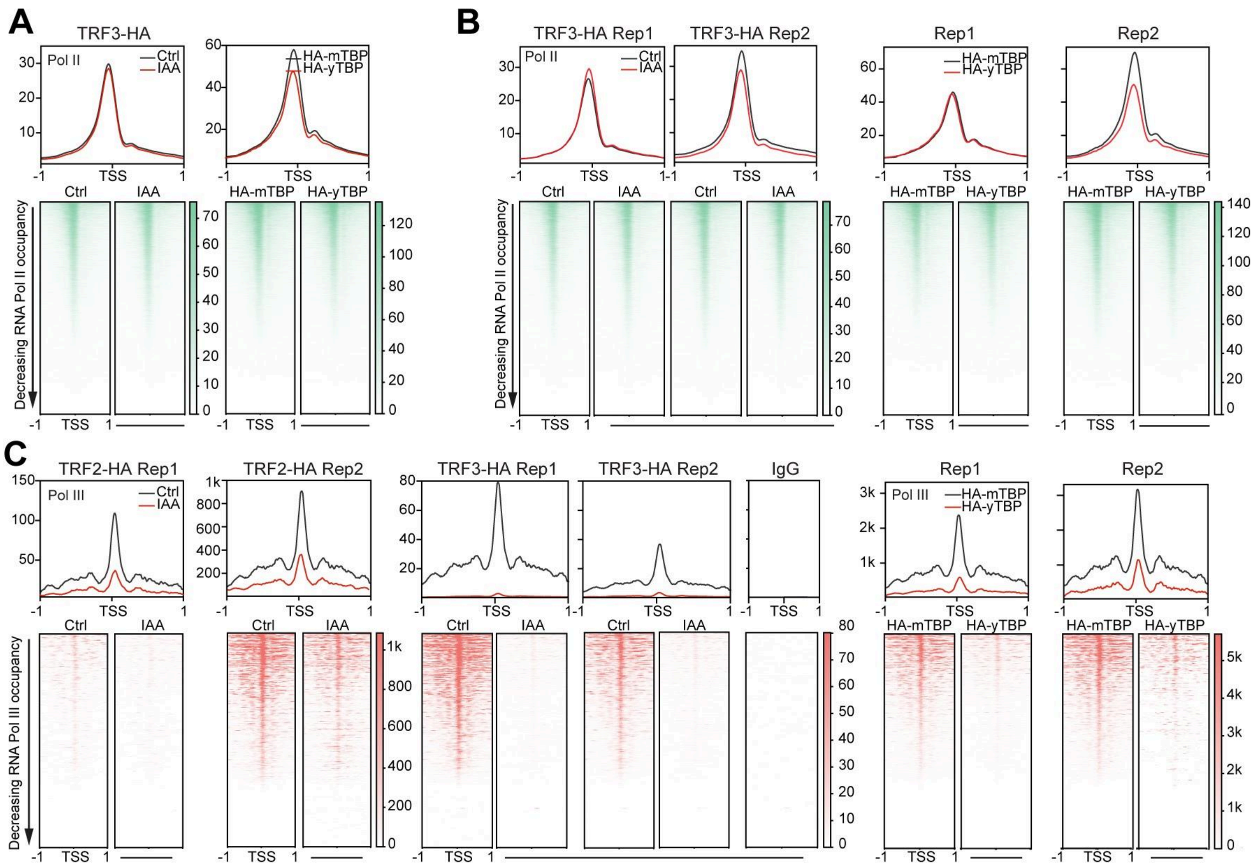


**Figure S3. Replicate analyses of RNA Pol II and III binding in the presence of TBP homologs**

(A) Genome-wide average plots (top) and heatmaps (bottom) arranged by decreasing Pol II occupancy of Pol II CUT&Tag in a 2kb window surrounding the TSS of all RNA Pol II genes. Samples shown are TRF3-HA in DMSO- (control) or IAA-treated conditions (left), HA-mTBP and HA-yTBP (right). (B) Biological replicates of (A). (C) Biological replicates of Pol III CUT&Tag analyses displayed as average plots (top) and heatmaps (bottom) in a 2-kb window centered at the TSS of all tRNA genes. Samples shown include TRF3-HA in DMSO- (control) and IAA-treated conditions, IgG as negative control for CUT&Tag signal, HA-mTBP, and HA-yTBP.

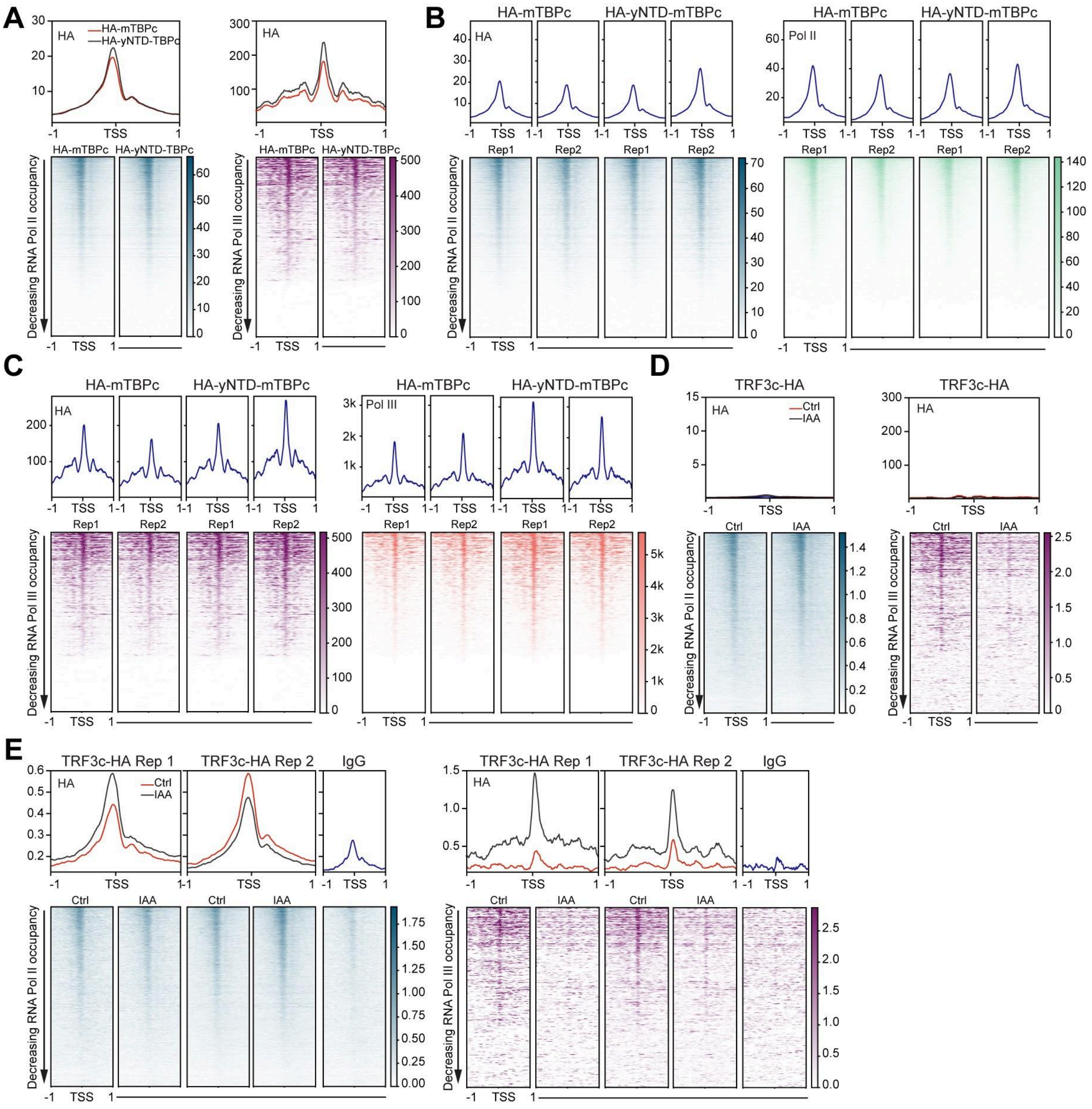


**Figure S4. Replicate analyses of NTD truncations and chimeric TBP proteins**

(A) Genome-wide average plots (top) and heatmaps (bottom) arranged by decreasing Pol II occupancy of HA CUT&Tag for HA-mTBPc and HA-yNTD-mTBPc in a 2 kb window surrounding the TSS of all RNA Pol II (left) and tRNA (right) genes. (B) Biological replicates of HA (left) and RNA Pol II (right) CUT&Tag analyses displayed as average plots (top) and heatmaps (bottom) in a 2-kb window centered at TSS of RNA Pol II genes. Samples include mESCs expressing HA-mTBPc and HA-yNTD-mTBPc. (C) Same as (B) but for all HA and

RNA Pol III CUT&Tag at tRNA genes. (D) Average plots (top) and heatmaps (bottom) for HA CUT&Tag in TRF3c-HA expressing mESCs are displayed for RNA Pol II (left) and tRNA (right) genes in control and IAA-treated conditions. (E) Biological replicates of (D) shown with IgG CUT&Tag control.

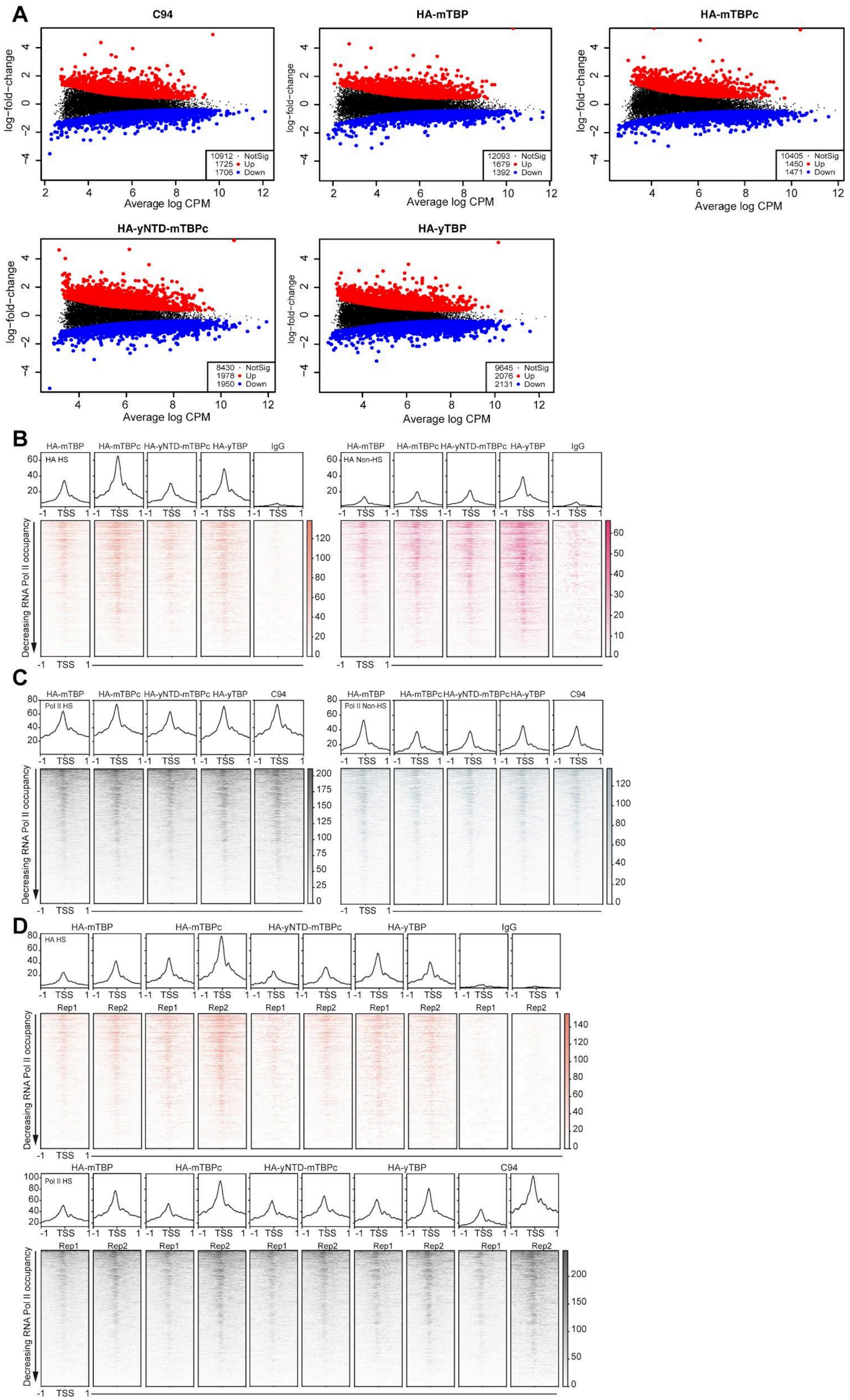
A
C94
HA-mTBP
HA-mTBPc
HA-yNTD-mTBPc
HA-yTBP
log−fold−change
Average log CPM
10912 · NotSig
1725 • Up
1706 • Down
12093 · NotSig
1679 • Up
1392 • Down
10405 · NotSig
1450 • Up
1471 • Down
8430 · NotSig
1978 • Up
1950 • Down
9645 · NotSig
2076 • Up
2131 • Down
B
HA-mTBP
HA-mTBPc
HA-yNTD-mTBPc
HA-yTBP
IgG
HA HS
HA Non-HS
TSS
Decreasing RNA Pol II occupancy
C
HA-mTBP
HA-mTBPc
HA-yNTD-mTBPc
HA-yTBP
C94
Pol II HS
Pol II Non-HS
TSS
Decreasing RNA Pol II occupancy
D
HA-mTBP
HA-mTBPc
HA-yNTD-mTBPc
HA-yTBP
IgG
HA HS
Rep1
Rep2
TSS
Decreasing RNA Pol II occupancy
HA-mTBP
HA-mTBPc
HA-yNTD-mTBPc
HA-yTBP
C94
Pol II HS
Rep1
Rep2
TSS
Decreasing RNA Pol II occupancy

**Figure S5. The role of NTD in transcription reprogramming during stress**

(A) Differential gene expression analyses of Pol II CUT&Tag in heat-shocked versus non-heat shock conditions for each HA-tagged homolog expressed in mESCs. The number of up- and down-regulated genes upon heat shock are indicated in red and blue, respectively. (B) Genome-wide average plots (top) and heatmaps (bottom) arranged by decreasing Pol II occupancy of HA CUT&Tag for HA-mTBP, HA-mTBPc, HA-yNTD-mTBPc, HA-yTBP and IgG as negative control in a 2 kb window surrounding the TSS of HS-induced genes at heat shock (left) or non-heat shock conditions (right). (C) Genome-wide average plots (top) and heatmaps (bottom) arranged by decreasing Pol II occupancy of Pol II CUT&Tag for HA-mTBP, HA-mTBPc, HA-yNTD-mTBPc, HA-yTBP and IgG as negative control in a 2 kb window surrounding the TSS of HS-induced genes at heat shock (left) or non-heat shock conditions (right). (D) Biological replicates of samples with heat shock conditions in (B) and (C).

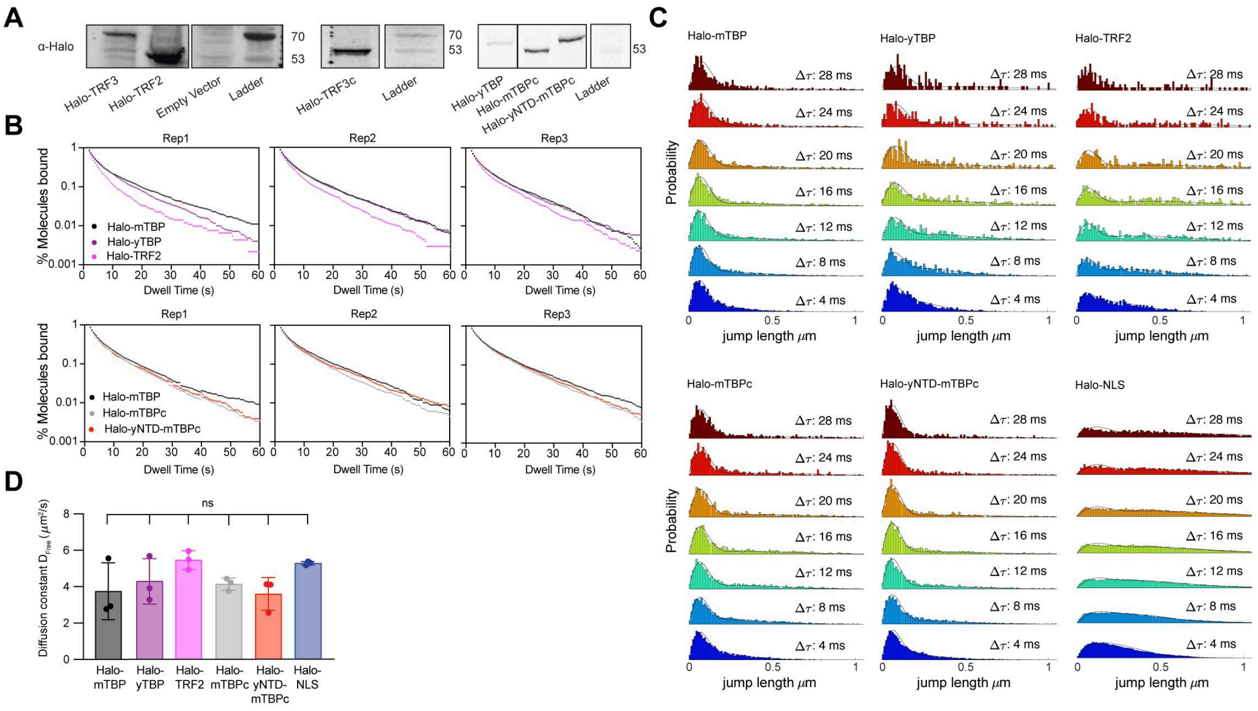


**Figure S6. Replicate analyses of TBP homolog dynamics**

(A) Western blot analyses of Halo-tagged homolog expressed in mESCs using α-Halo antibody. (B) Replicates of slow-tracking SPT. Dwell time histograms for each replicate are shown for each Halo-tagged homolog. (C) Jump length histograms were measured in displacements (μm) after 2-7 consecutive frames (Δτ = 4-28 ms) for each Halo-tagged homolog. (D) Diffusion coefficients of unbound molecules for each homolog extracted from fitting the jump length histogram with a 2-state kinetic model. Error bar represents SEM. Statistics: ns, non-significant.
